# Identification of INOSITOL PHOSPHORYLCERAMIDE SYNTHASE 2 (IPCS2) as a new rate-limiting component in Arabidopsis pathogen entry control

**DOI:** 10.1101/2025.01.09.632110

**Authors:** Josephine Mittendorf, Tegan M. Haslam, Cornelia Herrfurth, Nicolas Esnay, Yohann Boutté, Ivo Feussner, Volker Lipka

## Abstract

**SIGNIFICANCE STATEMENT:** Polarized transport of executive defense gene products to sites of attempted microbial invasion is important for plant pathogen entry control and disease resistance. Here, we provide evidence that INOSITOL PHOSPHORYLCERAMIDE SYNTHASE 2 (IPCS2)-dependent sphingolipid production contributes to the role of the trans-Golgi network as a multi-domain sorting compartment, and mediates proper delivery of an ATP-binding cassette transporter to polarized plasma membrane domains at plant-microbe interaction sites.

INOSITOL PHOSPHORYLCERAMIDE SYNTHASE 2 (IPCS2) is involved in the biosynthesis of complex sphingolipids at the *trans*-Golgi network (TGN). Here, we demonstrate a role of IPCS2 in penetration resistance against non-adapted powdery mildew fungi. A novel *ipcs2_W205*_* mutant was recovered from a forward genetic screen for Arabidopsis plants with enhanced epidermal cell entry success of the non-adapted barley fungus *Blumeria graminis* f. sp. *hordei* (*Bgh*). A yeast complementation assay and a sphingolipidomic approach revealed that the *ipcs2_W205*_* mutant represents a knock-out and lacks IPCS2-specific enzymatic activity. Further mutant analyses suggested that IPCS2-derived glycosyl inositol phosphorylceramides (GIPCs) are required for cell entry control of non-adapted fungal intruders. Confocal laser scanning microscopy (CLSM) studies indicated that upon pathogen attack, IPCS2 remains at the TGN to produce GIPCs, while focal accumulation of the defense cargo PENETRATION 3 (PEN3) at *Bgh* penetration sites was reduced in the *ipcs2_W205*_* mutant background. Thus, we propose a model in which sorting events at the TGN are facilitated by complex sphingolipids, regulating polar secretion of PEN3 to host-pathogen contact sites to terminate fungal ingress.

## INTRODUCTION

Plants have developed an efficient defense machinery to terminate the attack of non-adapted pathogens. As a first obstacle, pathogens encounter preformed physical and chemical barriers such as waxy cuticles, rigid cell walls and antimicrobial compounds (Thordal-Christensen 2003; Nürnberger and Lipka 2005). If intruders are able to successfully overcome these defense barriers, host cell surface-located pattern recognition receptors (PRRs) recognize conserved pathogen-associated molecular patterns (PAMPs) or host-derived danger-associated molecular patterns (DAMPs), triggering active defense responses (Jones and Dangl 2006; Boller and Felix 2009).

As obligate biotrophic pathogens, powdery mildew fungi derive nutrients from living host cells and cause disease on a wide range of plants, including economically important crops. During the asexual life cycle, conidiospores germinate on the plant leaf surface and produce appressoria to penetrate the epidermal cell wall. Upon fungal entry attempts, the host plant dynamically rearranges the cytoskeleton, aggregates organelles and secretes antifungal compounds to form defense structures called papillae underneath host-pathogen contact sites (Lipka et al. 2008; Hückelhoven and Panstruga 2011). Apoplastic papillae consist of callose, reactive oxygen species (ROS), phenolics and defense molecules, providing physical and chemical barriers for penetration resistance against fungal intruders (Thordal-Christensen et al. 1997; Ellis 2006; Underwood and Somerville 2008; Meyer et al. 2009). However, if the fungal pathogen is able to successfully breach the plant cell wall, haustoria are formed that are required for host nutrient uptake and effector secretion (Micali et al. 2008). As a host cell response, post-penetration mechanisms are activated, which typically include the generation of callose-enriched haustorial encasements and programmed cell death (PCD) of invaded cells (Lipka et al. 2005; Meyer et al. 2009).

PENETRATION 1 (PEN1), PEN2, PEN3 and PEN4 were identified in forward genetic screens as molecular components of Arabidopsis cell entry control against non-adapted powdery mildew fungi (Collins et al. 2003; Lipka et al. 2005; Stein et al. 2006; Hématy et al. 2020). PEN1 is a plasma membrane (PM)-resident SOLUBLE N-ETHYLMALEIMIDE-SENSITIVE-FACTOR ATTACHMENT RECEPTOR (SNARE) protein, which is required for vesicle fusion events to ensure papilla formation (Collins et al. 2003; Assaad et al. 2004). A second, PEN1-independent, pathway deals with the polarized secretion of antifungal compounds to sites of pathogen ingress. The mitochondria-associated atypical myrosinase PEN2 and the phytochelatin synthase PEN4 were reported to catalyze the generation of potentially toxic indole glucosinolate-derived molecules (Bednarek et al. 2009; Fuchs et al. 2016; Hématy et al. 2020). These antifungal products are believed to be secreted by the ATP binding cassette (ABC) transporter protein PEN3 across the PM to the apoplastic battlefield to terminate fungal penetration attempts (Stein et al. 2006; Bednarek et al. 2009; Hématy et al. 2020).

Interestingly, PM-recycled and not *de novo* synthesized PEN1 and PEN3 were found to be recruited to sites of attempted fungal invasion and strongly accumulate within apoplastic papillae (Meyer et al. 2009; Reichardt et al. 2011; Underwood and Somerville 2013). Further studies regarding the involvement of actin filaments revealed that polarized secretion of PEN1 and PEN3 follow distinct trafficking pathways (Underwood and Somerville 2013; Qin et al. 2021). However, the *trans*-Golgi-network (TGN)-based proteins aminophospholipid ATPase 3 (ALA3) and ECHIDNA (ECH) were reported to be required for delivery of both PEN1 and PEN3 to the cell surface (Uemura et al. 2019; Underwood et al. 2017; Liu et al. 2023). Thus, a model was suggested in which the receiving of recycled PEN1 and PEN3 is facilitated by a common TGN compartment, and from there splits into separate pathways for the polarized secretion of both defense proteins to sites of attempted fungal invasion (Nielsen 2024). In alignment with this hypothesis, the GDSL-lipase/esterase family protein GOLGI DEFECTS 36 (GOLD36), which alters the lipid composition at the TGN, was found to be a specialized component in the delivery of PEN3 to apoplastic papillae (Underwood 2022).

Upon infection with a non-adapted powdery mildew, the nanodomain-localized REMORIN 1.3 (REM1.3) protein accumulates in papillary membranes, providing evidence that these include some types of nanodomains (Xing et al. 2019). Further, PEN1 has been reported to co-localize with REM1.3 in papillae nanodomains (Xing et al. 2019). As nanodomains are important for membrane subcompartmentalization to act as, *inter alia*, signaling platforms in response to different stimuli (Liang et al., 2018; Platre et al., 2019; Smokvarska et al., 2020; Ma et al., 2022; Jaillais et al., 2024), they could here serve as transfer stations delivering immunity cargo such as PEN1 and PEN3 to extracellular papillae. Some plant membrane nanodomains are lipid-ordered and rely on sterol- and sphingolipid-composition (Grosjean et al., 2015; Gronnier et al., 2017; Lv et al., 2017). Although PEN3 was never shown to be part of nanodomains via co-localization studies using nanodomains marker proteins, biochemical analysis suggests that the membrane environment of PEN3 might be sterol- and sphingolipid-enriched (Minami et al. 2009; Kierszniowska et al. 2009). The most abundant sphingolipids at the PM are glycosyl inositol phosphorylceramides (GIPCs) (Bahammou et al., 2024). GIPCs contain a ceramide backbone, which consists of a saturated or monounsaturated very-long-chain fatty acid (VLCFA) and a poly-hydroxylated long-chain base (LCB) moiety, modified with a polar sugar headgroup (Luttgeharm et al. 2016; Mamode-Cassim et al., 2020; Haslam and Feussner 2022). The first committed step in GIPC production is catalyzed by inositol phosphorylceramide synthases (IPCS1-3), which transfer a head group from phosphatidylinositol (PI) onto ceramide to produce the intermediate inositol phosphorylceramide (IPC) at the TGN (Mina et al. 2010; Ito et al. 2021), which is then further modified by glycosylation events. Biophysical analyses revealed that GIPCs influence membrane properties by increasing thickness and electronegativity of the PM outer leaflet (Mamode Cassim et al. 2021). Moreover, previous studies provided evidence that IPCS2-derived GIPCs facilitate polar secretion of the auxin efflux carrier PIN2 to apical membranes, suggesting that GIPCs are involved in secretory trafficking (Markham et al. 2011; Wattelet-Boyer et al. 2016; Ito et al. 2021).

Here, we demonstrate a direct link between sphingolipid metabolism and Arabidopsis cell entry control. Genetic interference of *IPCS2* results in a defective penetration resistance against non-adapted powdery mildews. Mutant analyses combined with sphingolipidomics suggested that reduction in IPCS2-derived (G)IPCs causes a *pen* phenotype. Genetic analyses revealed that IPCS2 is involved in the PEN2/PEN3-mediated defense against fungal intruders. Furthermore, confocal laser scanning microscopy (CLSM) indicated that IPCS2-derived GIPCs at the TGN impact the polarized secretion of PEN3 to host-pathogen contact sites. Our results uncover a role of complex sphingolipids in polar secretory trafficking of the immunity cargo PEN3 to sites of attempted fungal invasion.

## RESULTS

### Mutations in *IPCS2* result in enhanced cell entry success of non-adapted powdery mildews

The *ipcs2_W205*_* mutant was isolated in a forward genetic screen for Arabidopsis plants that allow enhanced entry of the non-adapted barley fungus *Blumeria graminis* f.sp. *hordei* (*Bgh*) into epidermal pavement cells (Supplemental Figure 1). This mutation results in a premature stop codon, truncating IPCS2 at amino acid residue 205 (Figure 1A). As the truncation of IPCS2 results in the loss of one of two predicted catalytic motifs conserved among IPCSs (Wang et al., 2008; Huitema et al., 2004), the *ipcs2_W205*_* mutant likely represents a knock-out and lacks enzymatic activity of IPCS2 (Figure 1A). In the Arabidopsis ecotype Columbia 0 (Col-0), only 15% ± 4% of germinated *Bgh* conidiospores successfully penetrate the plant cell wall at 3 days post infection (dpi), whereas the remaining attempts are blocked by the formation of the defense structure papilla (Figure 1B). In contrast, the *Bgh* penetration frequency increases to 46% ± 10% on the leaves of *ipcs2_W205*_* plants (Figure 1B). To confirm that the mutation in *IPCS2* causes the enhanced cell entry success of *Bgh* in this mutant, we analyzed another *ipcs2* mutant, which has a transfer DNA (T-DNA) insertion 395 bp upstream of the translational start of *IPCS2* (Figure 1A). Like *ipcs2_W205*_*, *ipcs2*-1 was more susceptible than the wild-type Col-0 to *Bgh* penetration (Figure 1B). Repeating this pathogenicity assay with the non-adapted pea fungus *Erysiphe pisi* (*E.pisi*) yielded similar results, showing an increase of *E.pisi* entry rates up to a minimum of 45% ± 12% on both *ipcs2* mutants compared to the wild-type Col-0 (Figure 1C). Notably, the adapted powdery mildew *Golovinomyces orontii* (*G.orontii*) could grow on the *ipcs2* mutants at an equal level as on wild-type leaves (Figure 1D), suggesting that IPCS2 function in Arabidopsis cell entry control is restricted to non-adapted powdery mildews.

**Figure 1:**
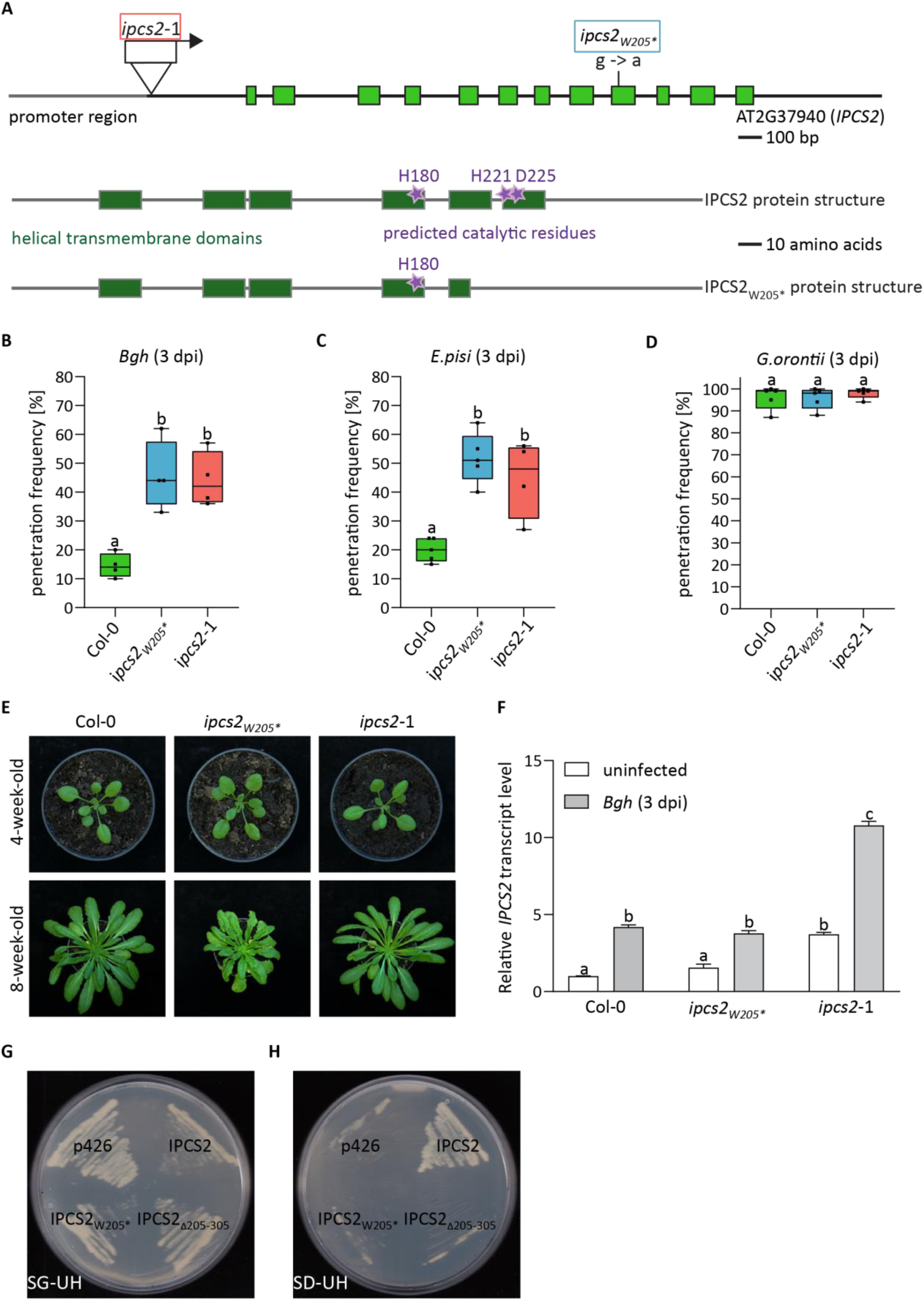
IPCS2 is involved in Arabidopsis cell entry control against non-adapted pathogens. A: Genomic organization of *IPCS2* including the promoter region and exon/intron structure, as well as the protein structure of IPCS2 and IPCS2_W205*_. The location of the ethyl methanesulfonate (EMS)-induced premature stop codon insertion in the *ipcs2_W205*_* mutant is marked with a blue box, while the red box indicates the location and orientation of the T-DNA insertion in the *ipcs2*-1 mutant. B-D: Penetration frequency of (B) *Bgh*, (C) *E.pisi* and (D) *G.orontii* on wild-type Col-0, *ipcs2_W205*_* and *ipcs2*-1 mutant leaves at 3 days post infection (dpi). Individual box plots include whiskers representing minimum and maximum values. Letters show significant differences between genotypes (n = 4-5; one-way ANOVA with Tukeýs post-hoc test; p<0.05). Similar results were confirmed in at least two additional independent experiments. E: Macroscopic phenotype of wild-type Col-0 and both *ipcs2* mutant variants after 4-or 8-weeks under short-day conditions. F: Uninfected (white bars) and *Bgh*-infected (3 dpi; grey bars) leaves of wild-type Col-0, *ipcs2_W205*_* and *ipcs2*-1 were analyzed by quantitative real-time PCR. Presented *IPCS2* transcript levels are relative to the uninfected Col-0 sample. The data represent the average ± standard error of the mean (n = 3-5). Letters show significant differences between genotypes and treatments (one-way ANOVA with Tukeýs post-hoc test; p < 0.05). The experiment was repeated three times with similar results. G-H: Complementation of yeast *YPH499-HIS-GAL1:AUR1* conditional mutant with the empty p426 vector (negative control), Arabidopsis wild-type *IPCS2*, *IPCS2_W205*_* and *IPCS2_Δ205-305_*. Expression of endogenous *AUR1* is induced in yeast cells grown on galactose-containing media (SG-UH; G) and suppressed on media supplemented with glucose instead of galactose (SD-UH; H). Images are representative of complementation (or non-complementation) in cells derived from four distinct colonies recovered from transformation.

Macroscopic analyses revealed that the *ipcs2_W205*_* mutant exhibits a decreased rosette size and a spontaneous lesion phenotype at later growth stages, while no pleiotropic phenotype is visible for the *ipcs2*-1 mutant (Figure 1E). To compare *IPCS2* transcript levels in both *ipcs2* mutants, we performed quantitative real-time PCR (qPCR) with uninfected and *Bgh*-infected (3 dpi) leaf samples of wild-type Col-0 and both *ipcs2* mutant alleles. Notably, this analysis revealed *Bgh*-induced expression of *IPCS2* in the wild-type Col-0 (Figure 1F). The *IPCS2* transcript levels in uninfected and challenged *ipcs2_W205*_* mutant plants were similar to the wild-type (Figure 1F), indicating that *IPCS2* transcript stability is not affected by the premature stop codon. In contrast, significantly elevated *IPCS2* expression was observed in uninfected *ipcs2*-1 leaves, and was even more upregulated upon pathogen attack (Figure 1F). Therefore, we concluded that the T-DNA insertion in the promoter region of *IPCS2* induces overexpression in the *ipcs2*-1 mutant, via a yet unknown mechanism.

As the premature stop codon insertion in *IPCS2* does not affect transcript stability, we determined whether the truncated IPCS2_W205*_ has residual enzymatic activity *in vivo* via a previously described yeast complementation assay (Mina et al. 2010). Thus, wild-type *IPCS2* (Mina et al. 2010), *ipcs2_W205*_* and the *ipcs2_Δ205-305_* variant, where the amino acids after the premature stop codon at position 205 are deleted, were cloned into a p426 expression vector (Sikorski and Hieter 1989; Mumberg et al. 1995) under the control of the strong, constitutive *Saccharomyces cerevisiae GLYCERALDEHYDE-3-PHOSPHATE DEHYDROGENASE* (*GPD*) promoter. These constructs were transformed into the conditional mutant *YPH499–HIS3– GAL1:AUR1* (Denny et al. 2006), in which a galactose-inducible promoter replaces the endogenous promoter of yeast *IPCS, AUREOBASIDIN A RESISTANCE1* (*AUR1*). Therefore, *AUR1* expression is induced by galactose-containing medium, while addition of glucose suppresses the expression. In contrast to the positive control wild-type IPCS2, both IPCS2_W205*_ mutant variants could not rescue growth of the *YPH499-HIS3-GAL1:AUR1* strain (Figure 1G, H), indicating that the truncated IPCS2_W205*_ protein has no residual enzymatic activity in yeast cells.

### Total ceramide content is enriched in *ipcs2_W205*_*, while total (G)IPC content is reduced

To further assess the effect of the *ipcs2_W205*_* mutation and the T-DNA insertion in *ipcs2*-1 *in planta*, we determined their sphingolipid profiles using targeted UPLC-nanoESI-MS/MS analysis. We measured ceramides, glucosylceramide (GlcCer), IPC and different classes of GIPCs. Briefly, the modification of ceramide with a glucose residue by glucosylceramide synthase (GCS) generates GlcCer (Leipelt et al. 2001; Melser et al. 2010) (Figure 2A). To produce GIPCs, IPCSs catalyze the addition of a head group derived from PI to ceramides for the production of the intermediate IPC (Mina et al. 2010) (Figure 2A). The next step involves the addition of a glucuronic acid (GlcA) moiety via an α-(1,4)-linkage to IPC, catalyzed by the inositol phosphorylceramide glucuronosyltransferase 1 (IPUT1) (Rennie et al. 2014) (Figure 2A). Further production of Hex-GIPC requires the GIPC mannosyl transferase 1 (GMT1) (Figure 2A), while HexNAc-GIPC synthesis depends on glucosamine inositol phosphorylceramide transferase 1 (GINT1) (Fang et al. 2016; Ishikawa et al. 2018). In comparison to the wild-type Col-0, ceramide levels were significantly enriched in *ipcs2_W205*_*, but not in *ipcs2*-1, whereas the GlcCer content was unaffected by the mutations in *IPCS2* (Figure 2B, C). Notably, IPC and Hex-GIPC levels were significantly reduced in *ipcs2_W205*_*, while *ipcs2*-1 showed similar levels as the wild-type Col-0 (Figure 2D, E). These results suggest that the premature stop codon insertion in *ipcs2_W205*_* leads to a knock-out mutation and loss of enzymatic activity of IPCS2, resulting in accumulation of ceramide and reduction in IPC as well as Hex-GIPC levels. In contrast, the sphingolipid profile of *ipcs2*-1 was wild-type-like, suggesting that the T-DNA insertion in the *IPCS2* promoter region does not impact the biochemical function of IPCS2 and sphingolipid homeostasis. Further investigations are necessary to clarify the underlying mechanism that causes the compromised penetration resistance in *ipcs2*-1. The remainder of the present study, however, is focused on the *ipcs2_W205*_* mutant.

**Figure 2:**
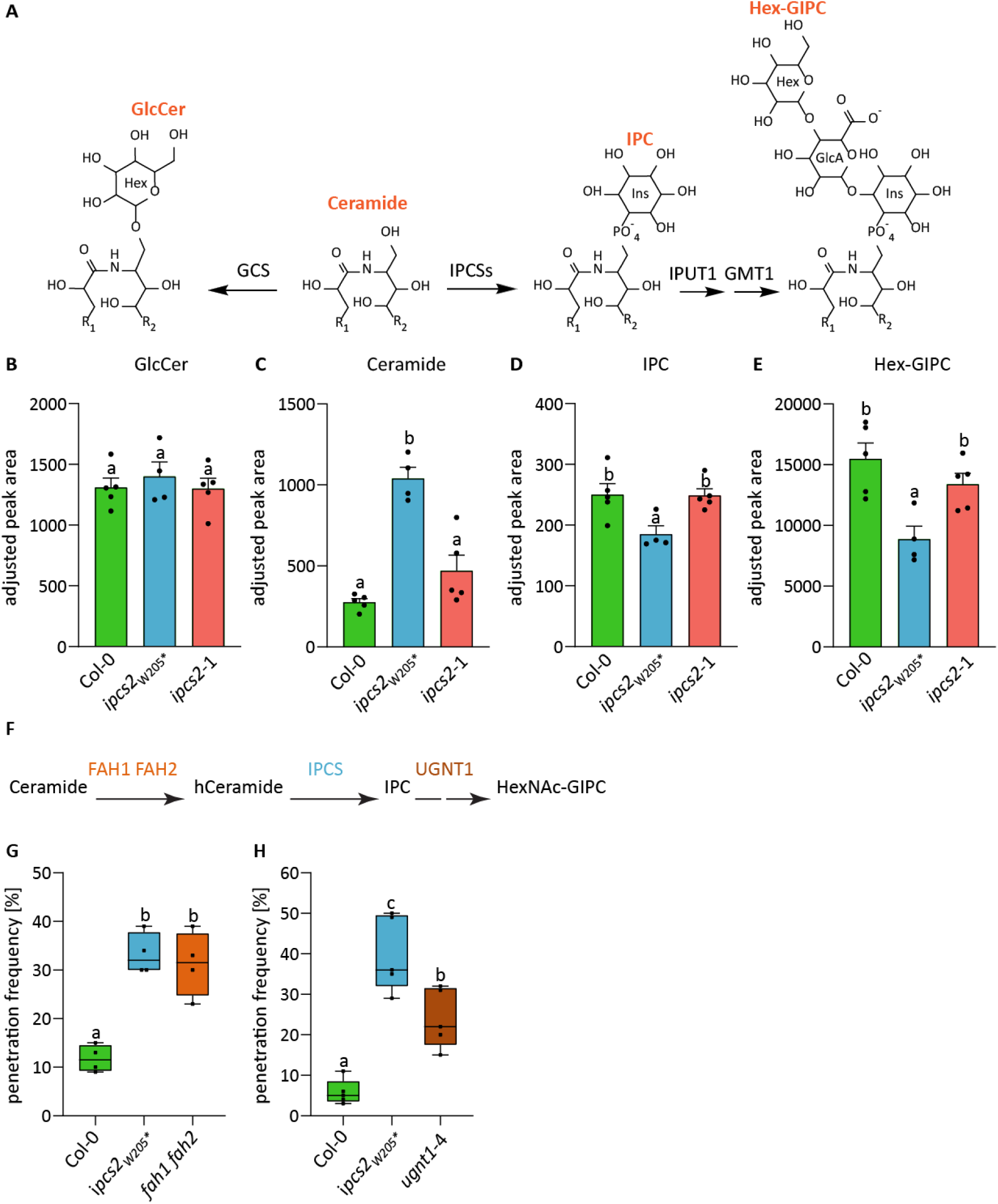
Sphingolipid profiles of Arabidopsis *ipcs2* mutants. A: Different metabolic fates of ceramide. Transfer of a glucose residue to ceramide produces GlcCer. To generate GIPCs, an inositol phosphate is transferred onto the ceramide backbone by IPCS. Subsequently, a GlcA moiety is added by IPUT1 and GMT1 catalyzes mannosylation of the head group. R_1_ = FA; R_2_ = LCB. B-E: Sphingolipid measurements of Arabidopsis rosettes for wild-type Col-0 and two *ipcs2* mutant alleles. Values are individual MRM peak areas normalized to the total fatty acid methyl ester (FAME) quantity and mg dry weight per sample. Bars represent averages of 4 to 5 biological replicates, error bars are standard error of the mean. Letters indicate significance at p < 0.05 determined by a one-way ANOVA with Tukey’s post-hoc test. Similar results were obtained in an independent experiment. F: The substrate production of IPCS2 is facilitated by FAH1 and FAH2, which hydroxylate ceramides. After transfer of inositol phosphate to hCeramide by IPCS, IPC can be further modified to Hex-GIPC or HexNAc-GIPC. The latter requires the activity of the UDP-GlcNAc transporter UGNT1. G-H: Box plots demonstrate the *Bgh* penetration frequency on wild-type Col-0, *ipcs2_W205*_* and (G) *fah1 fah2* or (H) *ugnt1*-4 leaves at 3 days post infection (dpi). Whiskers include minimum and maximum values. Letters show significant differences between genotypes (n = 4-5; one-way ANOVA with Tukeýs post-hoc test; p<0.05). Similar results were obtained in two additional independent experiments.

As the IPC and Hex-GIPC content levels in *ipcs2_W205*_* were significantly reduced but not fully abolished, we suspected that the IPCS2 homologs IPCS1 and IPCS3 may contribute to the synthesis of IPC in this mutant. Based on expression profiles, *IPCS2* is the most highly-expressed *IPCS* isoform in all tissues, while *IPCS1* exhibits weak constitutive expression (0.05% of *IPCS2* levels in rosette leaves) (Mina et al. 2010). In contrast, *IPCS3* has high constitutive expression in stems and flowers, and is inferred to have a specialized role in these tissues (Mina et al. 2010). As a common problem with strong alteration of GIPC levels, double knockout *ipcs1 ipcs2* mutants could not be analyzed due to seedling lethality (Tartaglio et al. 2017; Ito et al. 2021). Therefore, we analyzed the previously described Arabidopsis inducible *IPCS1;2^amiRNA^* knockdown line (Ito et al. 2021). In contrast to the *ipcs2_W205*_* mutant, the *Bgh* penetration frequency was only slightly increased for the *IPCS1;2^amiRNA^* knockdown line after β-Estradiol (E2) treatment, but not significantly different from the wild-type Col-0 (Supplemental Figure 2A). This attenuated effect might be due to the E2-induced decrease in *IPCS1* and *IPCS2* transcript level of only 54% and 48%, respectively, in the *IPCS1;2^amiRNA^* line (Supplemental Figure 2B, C). This relatively mild phenotype compared to the *ipcs2_W205*_* mutant allele suggests that IPCS2, and not IPCS1, has a predominant role in cell entry control. This was further supported by the finding that the *ipcs1* single knock-out exhibits a wild-type-like *Bgh* infection phenotype (Supplemental Figure 2D). Interestingly, similar to *IPCS2*, *IPCS1* transcript levels were increased by *Bgh* infection in the wild-type Col-0 and were even more upregulated in the *ipcs2_W205*_* mutant (Supplemental Figure 2E). Taken together, our results suggest that IPCS2 is the major contributor to cell entry control, while in the absence of IPCS2 *IPCS1* is likely upregulated to partially complement this function.

### IPCS2-derived IPC and Hex(NAc)-GIPCs are required for Arabidopsis pathogen entry control

To further assess whether the reduction of IPC and Hex-GIPC levels in *ipcs2_W205*_* cause the enhanced cell entry success of non-adapted powdery mildews, we tested the double mutant of FATTY ACID HYDROXYLASE 1 and 2 (FAH1/2), which hydroxylate the α position of the fatty acid moiety in ceramides, thereby contributing to the production of the IPCS2 substrate (König et al. 2012; Nagano et al., 2012; König et al. 2022) (Figure 2F). Similar to *ipcs2_W205*_*, the penetration frequency of *Bgh* was increased up to 31% ± 6% on *fah1 fah2* leaves compared to the wild-type Col-0 (Figure 2G). This supports the hypothesis that the products of IPCS2 are required for cell entry control, rather than a moonlighting function of the IPCS enzyme. To determine whether other types of GIPCs than Hex-GIPCs, which are the predominant GIPC form in Arabidopsis, contribute to cell entry success of non-adapted powdery mildews, we examined knock-out mutants of *UDP-GlcNAc TRANSPORTER 1* (*UGNT1*), deficient in a sugar nucleotide transporter required for production of HexNAc-GIPCs (Ebert et al. 2018) (Figure 2F). As HexNAc-GIPC levels were too low for detection by LC-MS, this genetic approach was necessary. Mutant analysis of *ugnt1*-4 (Figure 2F) revealed elevated cell entry success rates of *Bgh* (Figure 2H). However, the effect was not as drastic as genetic interference in *ipcs2* and *fah1 fah2*. Thus, not only IPCS2-derived Hex-GIPCs, but also HexNAc-GIPCs could support penetration resistance against non-adapted powdery mildew fungi.

### IPCS2 is involved in the Arabidopsis PEN2/PEN3-dependent pathogen entry control pathway

In steady-state, IPCS2 was found to co-localize with the SYNTAXIN OF PLANTS 61-secretory vesicle (SYP61-SV) subdomain of the TGN (Ito et al. 2021). The next step was to track the subcellular behavior of IPCS2 and the associated (G)IPC production upon pathogen attack. Therefore, stable transgenic Col-0 and *ipcs2_W205*_* plants expressing *pUBQ:IPCS2-mVenus* were generated. Infection phenotype analyses confirmed that IPCS2-mVenus can restore the *Bgh* penetration rates on *ipcs2_W205*_* mutant leaves to wild-type levels (Supplemental Figure 3A). CLSM studies revealed that this functional IPCS2-mVenus fusion protein does not accumulate at *Bgh* infection sites in neither wild-type nor *ipcs2_W205*_* mutant plants (Figure 3A). This suggests that IPCS2 is not subjected to pathogen-induced polarization to infection sites, and rather stays at the TGN for (G)IPC production during pathogen attack.

**Figure 3:**
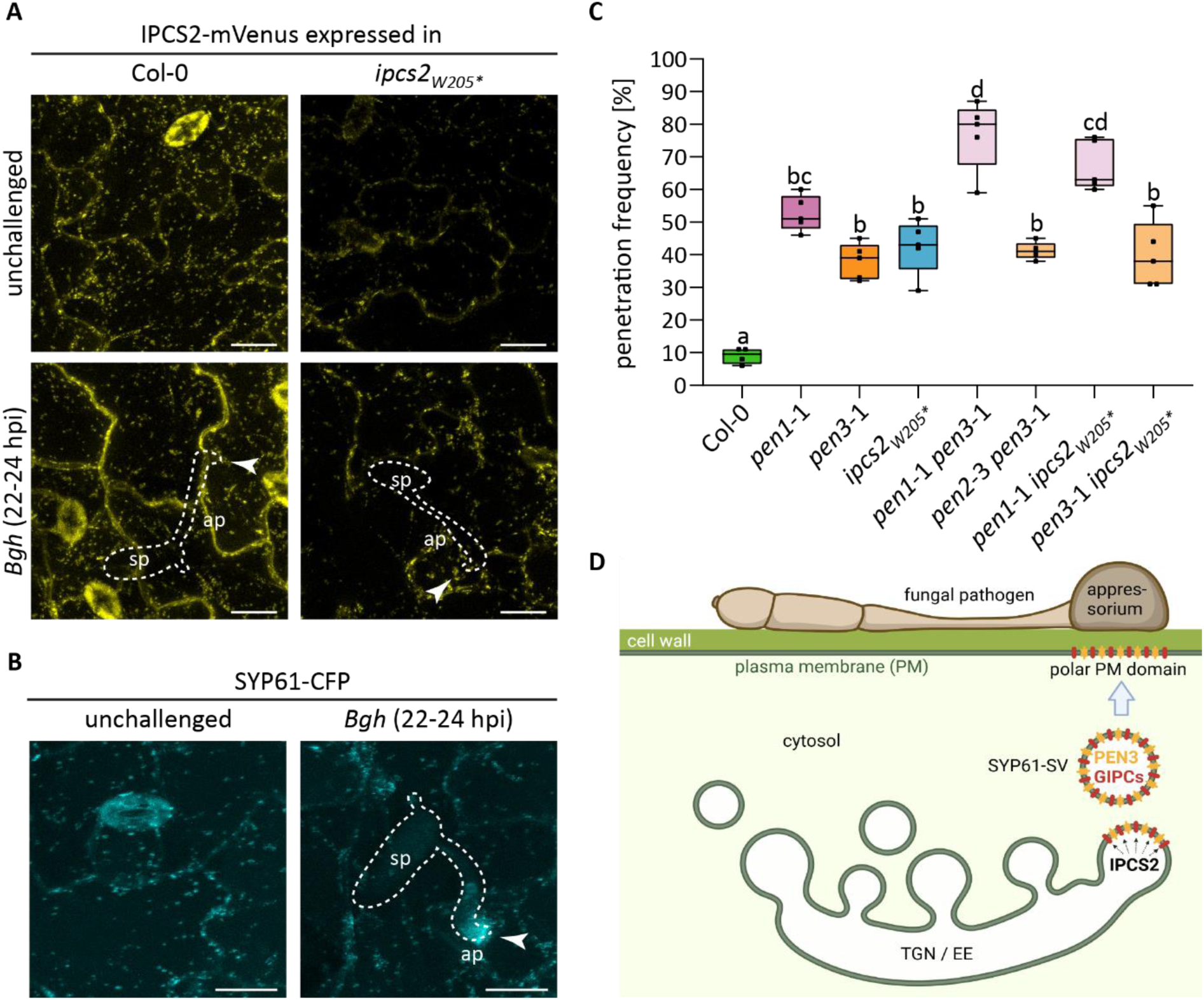
IPCS2 is part of the PEN2/PEN3 pathway. A: Stable transgenic Col-0 and *ipcs2_W205*_* plants expressing *pUBQ10:IPCS2-mVenus* were analyzed by CLSM. Maximum z-projections of CLSM images from unchallenged or *Bgh*-infected (22-24 hours post infection (hpi)) leaf epidermal cells are presented. *Bgh*-plant interaction sites are marked with a white arrowhead. White lines indicate the fungal spore with appressorium formation. Ap = appressorium; sp = spore. Scale Bar = 20 µm. B: Maximum z-projections of CLSM images from unchallenged or *Bgh*-infected (22-24 hpi) leaf epidermal cells of Col-0 plants stably expressing *pSYP61:SYP61-CFP*. Scale bar = 10 µm. C: Box plots illustrate the *Bgh* penetration frequency on the indicated single and double mutants at 3 days post infection. Whiskers represent minimum and maximum values, while letters show significant differences between genotypes (n = 5; one-way ANOVA with Tukeýs post-hoc test; p<0.05). D: Model illustrating IPCS2-dependent production of GIPCs, which are incorporated into SYP61-SVs and likely shuttle defense cargo such as PEN3 to polar domains underneath sites of attempted fungal invasion. Model created with Biorender.com. All results shown in this figure were replicated in two additional independent experiments.

Emerging evidence suggests that SYP61-SVs play a crucial role in the polar transport of GIPC-enriched cargo from the TGN to polar domains of the PM (Tapken and Murphy 2015; Wattelet-Boyer et al. 2016). CLSM analysis of a transgenic SYP61-CFP marker line revealed focal accumulation of SYP61-CFP at the tip of the *Bgh* appressorium (Figure 3B), indicating that SYP61-SVs are involved in the polarized secretion of defense material to domains underneath sites of fungal invasion attempts. As previous proteomic studies of the SYP61-SVs/TGN immuno-purified fractions revealed PEN1 and PEN3 as cargo of SYP61-SVs (Ito et al. 2021), we hypothesized that IPCS2-derived GIPCs could facilitate the shuttling of PEN1 and/or PEN3 to polar domains at sites of attempted fungal invasion. As genetic analyses revealed that PEN1 and PEN2/PEN3 act in different pathways (Lipka et al. 2005; Stein et al. 2006), we performed the same approach with *ipcs2_W205*_* mutants to clarify in which pathway GIPCs are involved. To generate double mutants, *ipcs2_W205*_* was crossed to *pen1*-1 and *pen3*-1 single mutants. The *Bgh* penetration rate of the control *pen1*-1 *pen3*-1 double mutant was elevated in comparison to the single mutants, while the cell entry success of *Bgh* on *pen2*-3 *pen3*-1 leaves was similar to the single mutants (Figure 3C), confirming previous findings (Stein et al. 2006; Johansson et al. 2014). In contrast to *pen3*-1 *ipcs2_W205*_* double mutants, the *Bgh* penetration rate was increased for *pen1*-1 *ipcs2_W205*_* in comparison to the single mutants (Figure 3C), indicating that IPCS2 is more likely involved in the PEN2/PEN3-dependent pathway. Thus, we propose a model in which IPCS2 strongly produces (G)IPCs at the TGN upon powdery mildew attack (Figure 3D). These GIPCs are incorporated into SYP61-SVs for shuttling PEN3 to polar domains at sites of attempted fungal invasion (Figure 3D). GIPCs were previously reported to mediate PI4P consumption at the TGN and subsequent polar sorting of the auxin efflux carrier PIN2, via activation of phosphoinositide-specific phospholipases C (PI-PLC) (Ito et al. 2021). We hypothesized that PI4P consumption could also be a mechanism of IPCS2-mediated polar recruitment of immunity cargo such as PEN3 to sites of attempted fungal invasion. Upon infection with *Bgh*, PI4P accumulated at the tip of the appressorium (Supplemental Figure 4). However, as PI4P signals were extremely weak in leaf tissue, we failed to correlate any change in PI4P distribution at *Bgh* penetration sites with loss of IPCS2 function in the *ipcs2_W205*_* mutant background.

### Focal accumulation of PEN3 at sites of attempted fungal invasion requires IPCS2 activity

To test the hypothesis that IPCS2-derived GIPCs are required for polar transport of PEN3 to sites of attempted fungal invasion, we crossed *ipcs2_W205*_* with the published transgenic protein marker line *pPEN3:PEN3-GFP* (Stein et al. 2006). In CLSM analyses, PEN3-GFP fluorescence intensity of focal accumulation (FA) patterns at *Bgh* appressoria appeared to be reduced in the *ipcs2_W205*_* mutant background compared to the mother plant at 24 hours post infection (hpi) (Figure 4A). To evaluate whether this reduction of PEN3 FA at *Bgh* penetration sites is caused by delayed recruitment, we performed a time course experiment. Therefore, the frequency of PEN3-GFP FA patterns at *Bgh* penetration sites was assessed at 12 hpi and 24 hpi. The frequency of PEN3-GFP FA at *Bgh* appressoria was similar between the control line and PEN3-GFP in *ipcs2_W205*_* at both time points (Figure 4B). Thus, we hypothesized that dysfunction of IPCS2 enzymatic activity could impair the spatial rather than temporal recruitment of PEN3-GFP to sites of attempted fungal invasion, or perhaps the extent and intensity of PEN3 aggregation. To test this idea, the area and mean intensity of these PEN3-GFP FA patterns in the mother plant and the *ipcs2_W205*_* mutant background were quantified at 24 hpi. This quantification analysis demonstrated that the area of PEN3-GFP FA patterns at *Bgh* appressoria is unaffected by IPCS2 dysfunction, whereas the fluorescence intensity was significantly reduced in the *ipcs2_W205*_* mutant by 53% ± 11% relative to the mother plant (Figure 4C, D). In contrast, the recruitment of callose as a molecular marker of papillae was unaffected in the *ipcs2_W205*_* mutant (Supplemental Figure 5). Taken together, these results suggest that IPCS2 enzymatic activity is involved in the specialized delivery of PEN3 to host-powdery mildew contact sites.

**Figure 4:**
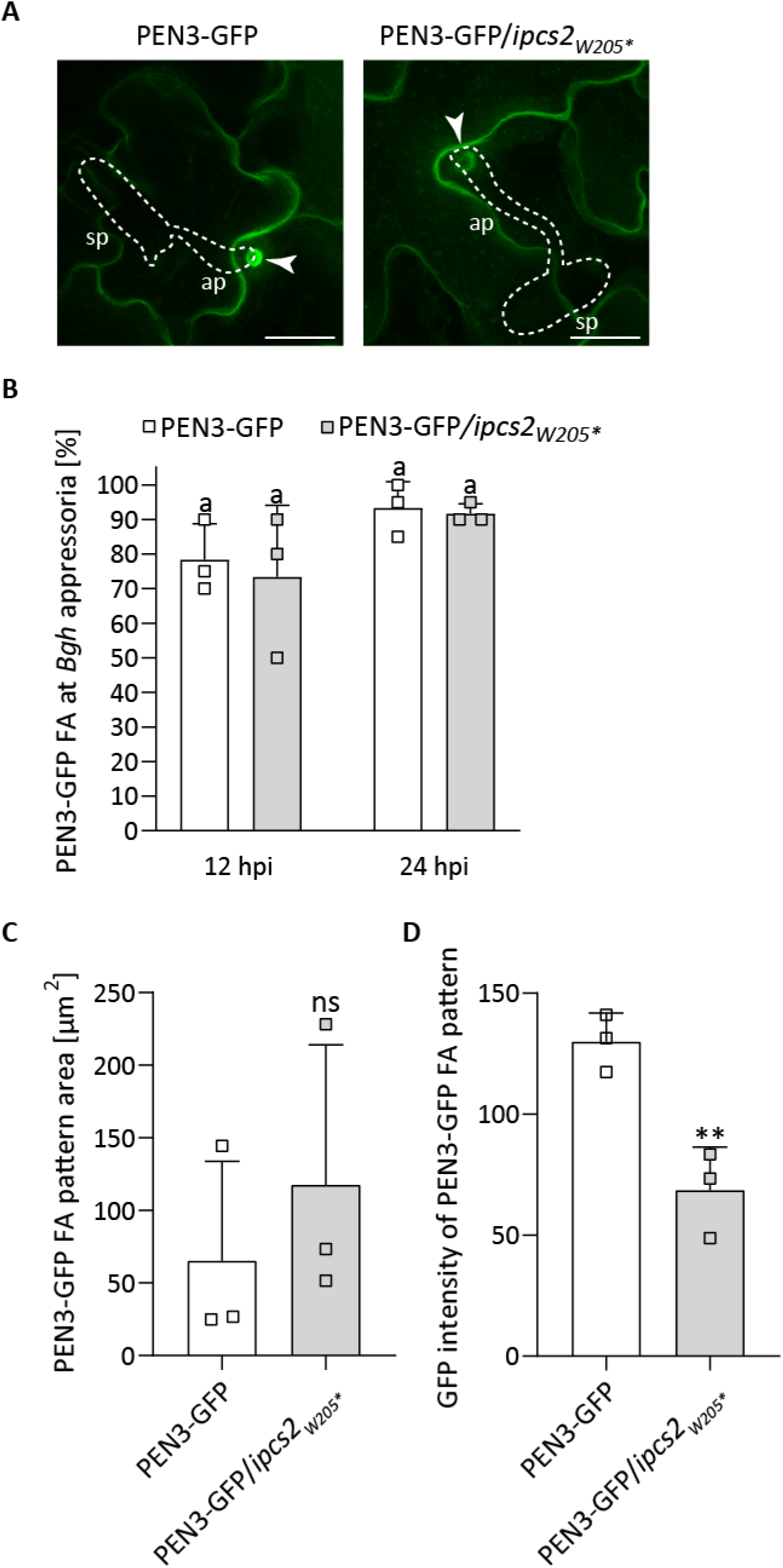
IPCS2 enzyme activity plays a crucial role in the polarized secretion of PEN3 to sites of attempted fungal invasion. A: Maximum z-projections of CLSM images reveal focal accumulation (FA) of PEN3-GFP at *Bgh* (24 hours post infection (hpi)) penetration sites (arrowheads) in the mother plant (left panel) or in the *ipcs2_W205*_* mutant background (right panel). White lines indicate the fungal spore with appressorium formation. Ap = appressorium; sp = spore. Scale Bar = 20 µm. B: Time-course evaluation of PEN3-GFP FA at plant-*Bgh* contact sites at 12 hpi and 24 hpi in the mother plant (white bars) or in the *ipcs2_W205*_* mutant background (grey bars). PEN3-GFP FA patterns were scored at the tip of 20 *Bgh* appressoria. Bars represent averages of three biological replicates ± standard deviation. Letters indicate significance at p < 0.05 determined by one-way ANOVA with Tukey’s post-hoc test. C-D: Quantification of (C) the area [µm^2^] or (D) the GFP intensity of PEN3-GFP FA patterns at the tip of 17-20 *Bgh* appressoria (24 hpi) in the mother plant (white bars) or in the *ipcs2_W205*_* mutant background (grey bars). Bars show the average and error bars represent standard deviation (n = 3). Significant differences to the mother plant were calculated with a Student’s *t*-test (ns = non-significant; ** = p < 0.0077). All experiments shown in this figure were repeated three times with similar outcomes.

## DISCUSSION

IPCS2 is primarily known for its involvement in the synthesis of complex sphingolipids (Mina et al. 2010). Here, we demonstrate that genetic interference of *IPCS2* causes enhanced cell entry success of non adapted powdery mildews, indicating a role of IPCS2 in penetration resistance. In this study, the novel *ipcs2_W205*_* mutant was identified, in which a premature stop codon results in loss of enzymatic activity in yeast cells and *in planta*. Our sphingolipidomic analysis revealed that compromised IPCS2 function results in ceramide accumulation and reduction of (G)IPC content, confirming published findings (Ito et al. 2021). *IPCS2* was previously recovered from an enhancer screen for RESISTANCE TO POWDERY MILDEW 8 (RPW8), which is involved in post-penetration resistance to several powdery mildew species (Xiao et al. 2001; Wang et al. 2008). Notably, the study of Wang et al. (2008) demonstrated that increased ceramide levels in the absence of IPCS2 induce salicylic acid (SA) accumulation, resulting in a lesion phenotype. In alignment with these findings, the *ipcs2_W205*_* mutant exhibited spontaneous cell death at later growth stages, which was abolished when SA synthesis (SALICYLIC ACID INDUCTION DEFICIENT 2 (SID2)) and signaling pathways were blocked (ENHANCED DISEASE SUSCEPTIBILITY 1, EDS1; PHYTOALEXIN-DEFICIENT, PAD4) by crossing the *ipcs2_W205*_* mutant to the triple *eds1*-2/*pad4*-1/*sid2*-2 (*eps*) mutant (Zhang et al. 2018) (Supplemental Figure 6). Furthermore, constitutive expression of IPCS2-mVenus under the strong *UBQ10* promoter in Col-0 and in the *ipcs2_W205*_* mutant resulted in spontaneous cell death (Supplemental Figure 3B), indicating that not only disruption but also overexpression of IPCS2 causes lesions. This supports the current model based on several publications (Wang et al. 2008; König et al. 2012; Bi et al. 2014; Simanshu et al. 2014; Zeng et al. 2021; Zeng and Yao 2022) that normal and controlled sphingolipid metabolism plays a crucial role in SA homeostasis, and that ceramides act as signaling lipids that induce PCD via SA. Thus, IPCS2 is not only required for cell entry control, but also for regulating ceramide levels to trigger post-penetration defense mechanisms.

In addition to the *ipcs2_W205*_* mutant, we analyzed another *ipcs2* mutant with defective penetration resistance, *ipcs2-*1, which has a T-DNA insertion at the end of the promoter region of *IPCS2*. qPCR revealed that *IPCS2* transcript levels are upregulated in uninfected *ipcs2*-1 plants, suggesting that the T-DNA insertion induces an increase in expression either by disrupting negative regulatory sequences or by insertion of activating sequences encoded by the T-DNA. However, *ipcs2*-1 mutants exhibit a wild-type-like macroscopic phenotype, while Col-0 plants stably expressing IPCS2-mVenus from a strong constitutive promoter showed a lesion phenotype (Supplemental Figure 3B). This suggests that IPCS2 protein may not constitutively hyperaccumulate in the *ipcs2*-1 mutant, which is supported by a wild-type-like sphingolipid profile. This is in alignment with the findings of a previous study reporting only subtle differences between wild-type Col-0 and *ipcs2*-1 in a sphingolipidomic analysis (Yan et al. 2019). However, the qPCR and sphingolipidomic data are based on a single leaf or whole rosette, respectively. Therefore, different cell type-specific expression of *IPCS2*, cell type-specific activity changes, or different subcellular behavior upon pathogen attack cannot be ruled out. As powdery mildew infection is restricted to the epidermis (Hückelhoven and Panstruga 2011), it would be interesting to analyze the epidermal expression profile of *IPCS2* in this T-DNA insertional mutant. Furthermore, *ipcs2_W205*_* mutant plants could be generated that stably express native IPCS2 tagged to mVenus from the *ipcs2*-1 promoter. This would allow subcellular tracking of the IPCS2 mutant variant in the epidermis upon pathogen attack.

Furthermore, *IPCS2* transcript levels were found to be elevated upon *Bgh* infection, confirming previous findings showing that *IPCS2* is upregulated by non-adapted powdery mildews (Nishimura et al. 2003; Stein et al. 2006; Wang et al. 2008). This pathogen-induced transcription was also observed in the *ipcs2*-1 mutant. In transgenic lines stably expressing IPCS2-mVenus under the control of a constitutive promoter in the wild-type or *ipcs2_W205*_* mutant background, IPCS2-mVenus protein levels were increased upon *Bgh* infection in Western blot analyses (Supplemental Figure 3C). This suggests a post-transcriptional stabilization of *IPCS2* RNA and/or IPCS2 protein upon pathogen attack, which presumably serves to increase GIPC production to terminate fungal ingress.

Mutant analyses of other components functioning upstream or downstream of IPCS2 lend further support to the idea that IPCS2-derived GIPCs are required for cell entry control against non-adapted fungal pathogens. Previous findings provided evidence that GIPCs contribute to the polar sorting of the auxin efflux transporter PIN2 without affecting endocytosis or recycling (Markham et al. 2011; Wattelet-Boyer et al. 2016). Here, we demonstrate that IPCS2-derived GIPCs are also required for polarized secretion of PEN3 to sites of attempted fungal invasion. In contrast, genetic analyses excluded IPCS2 function from the PEN1-dependent pathway and time-course experiments revealed that callose deposition at papillae is IPCS2-independent. As polar transport of the callose synthase POWDERY MILDEW RESISTANT 4 (PMR4)/GLUCAN SYNTHASE-LIKE 5 (GSL5) from the TGN to sites of attempted fungal invasion presumably depends on the PEN1 pathway (Assaad et al. 2004; Nielsen et al. 2012; Qin et al. 2021; Rubiato et al. 2022), we infer that IPCS2 acts in a specialized polar trafficking pathway for PEN3 recruitment. The transport model recently proposed by Nielsen (2024) suggests that recycled PEN1 and PEN3 overlap at the TGN upon pathogen infection, which is followed by splitting into distinct pathways for recruitment of both PEN proteins to sites of attempted fungal invasion. As PEN1, PMR4, PEN3 and IPCS2 were reported to be localized to the SYP61-SV subdomain of the TGN (Drakakaki et al. 2012; Ito et al. 2021), it is likely that this represents the shared TGN compartment after recycling. In this study, we demonstrate that upon pathogen attack, IPCS2-mVenus remains at the TGN, while SYP61-CFP accumulates at sites of attempted fungal invasion. Thus, IPCS2-dependent sorting processes of PEN3 take place at the TGN, potentially followed by transport via SYP61-tagged vesicles to fungal ingress sites. IPCS2-derived GIPC association with other lipids or proteins could potentially direct PEN3 as a specific cargo for transport to pathogen-induced polar domains, while PEN1 is excluded and follows a sphingolipid-independent secretory mechanism. Taken together, we provide evidence for a mechanistic link between IPCS2-derived complex sphingolipids and proper sorting of PEN3 to sites of attempted fungal invasion, uncovering a GIPC-dependent polar secretory trafficking pathway in Arabidopsis cell entry control.

## MATERIALS AND METHODS

### Plant material and growth conditions

The *Arabidopsis thaliana* ecotype Columbia (Col-0) was used as the wild-type. The following previously characterized single/double/triple mutants were analyzed: *ipcs1* (Ito et al. 2021), *fah1 fah2* (König et al. 2022), *pen1*-1 (Collins et al. 2003), *pen3*-1, *pen1*-1 *pen3*-1, *pen2*-3 *pen3*-1 (Stein et al. 2006) and *eps* (*eds1*-2/*pad4*-1/*sid2*-2) (Zhang et al. 2018). The seeds of the T-DNA insertion mutants *ipcs2*-1 (SALK_206784) and *ugnt1*-4 (SALK_075727) were obtained from the Nottingham Arabidopsis Stock Centre and genomic insertions of the T-DNAs were confirmed by PCR. All primers used in this study are listed in Supplemental Table S1. Double and quadruple mutants of *ipcs2_W205*_* were generated by crossing with the respective single/triple mutant and homozygous mutants were confirmed by PCR and Sanger sequencing. The following previously described Arabidopsis transgenic fluorescent protein marker lines were used: *pSYP61:SYP61-CFP* (Robert et al. 2008), *pPEN3:PEN3-GFP* (Stein et al. 2006) and *pUBQ10:mCITRINE-3×PH^FAPP1^* (Simon et al. 2014). Additionally, the recently established β-Estradiol-inducible *IPCS1;2* artificial microRNA line (Ito et al. 2021) was used for a pathogenicity assay.

Arabidopsis plants were grown under short-day conditions (8-h photoperiod, 22°C (day)/18°C (night), 65% relative humidity and a light intensity of approximately 150 μmol m^−2^ s^−1^) and used for experiments after 4 weeks.

### Generation of a pUBQ10:IPCS2-mVenus line

The previously described *pUBQ10:IPCS2-mVenus* construct was designed using the multisite gateway system (Nicolas Esnay 2020). The construct was confirmed by Sanger sequencing, transformed into *Agrobacterium tumefaciens* C1C58 and used for stable transformation of Col-0 and *ipcs2_W205*_* plants according to the floral dip method (Clough and Bent 1998). To select transgenic plants, soil-grown T1 seedlings were sprayed with BASTA (Bayer CropScience).

### Arabidopsis inoculation and powdery mildew pathogenicity assay

*Blumeria graminis* f.sp. *hordei* isolate K1 conidiospores were generated on *Hordeum vulgare* cv Lottie for 10 to 14 days, *Erysiphe pisi* (*E.pisi*) spores were produced on *Pisum sativum* (kleine Rheinländerin) for 14 days and *Golovinomyces orontii* (*G.orontii*) was cultivated on 4-week-old Arabidopsis plants for 10-14 days. For pathogenicity assays, 4-week-old Arabidopsis plants were infected with *Bgh*, *E.pisi* and *G.orontii*. After 22-24 hpi plants were used for CLSM and after 3 dpi for screening and quantification of fungal invasive growth rates. Visualization of papillae and haustoria formation as well as epidermal cell death was performed according to the established method by Lipka et al. (2005). For calculation of the penetration frequency (number of haustoria + epidermal cell death), 100 interaction sites per leaf of 3-5 individual plants per genotype were scored.

### Arabidopsis *pen* mutant screen and mapping of *ipcs2_W205*_*

M6 populations of Col-0 homozygous EMS mutant (HEM) lines (Capilla-Perez et al. 2018) were obtained from the Versailles Arabidopsis stock center. Plants were inoculated with *Bgh* and 3 days later screened for enhanced autofluorescence, indicative of cell death due to fungal entry, using a GFP1 filter set (excitation filter 425/60 nm; dichroic mirror 480 nm; Leica Microsystems) on a Leica M165 FC fluorescence-stereomicroscope. For mapping, a previously reported approach (Hartwig et al. 2012) was adapted. Thus, the *ipcs2_W205*_* mutant was backcrossed to Col-0. Approximately 380 F2 plants were screened for enhanced cell death rates. 28 plants with a clear *pen* phenotype were pooled and subjected to 150-bp paired end Illumina sequencing. The resulting reads were mapped to the Col-0 TAIR10 genome by using the CLC genomics workbench software and single nucleotide polymorphisms (SNPs) with a frequency of 20% or higher were called. SNPs present in Col-0 were subtracted from SNPs found in *ipcs2_W205*_*. The frequency of the remaining SNPs was plotted against their chromosomal position. The observed frequency peak on chromosome 2 contained a SNP resulting in a premature stop codon (W205*) in *IPCS2* (AT2G37940) (Supplemental Figure 1). A complementation analysis confirmed this polymorphism as the causative mutation.

### Inhibitor treatment

For amiRNA induction, 4-week-old plants of the *IPCS1;2^amiRNA^* line as well as the control plants were sprayed with 25 µM β-Estradiol (Sigma-Aldrich) and used for infection experiments after 48 h.

### Quantitative RT-PCR

RNA was extracted from Arabidopsis leaves using the innuPREP Plant RNA Kit (Innuscreen). cDNA was generated with RevertAid H Minus m-MuLV Reverse Transcriptase (Thermo Fisher Scientific). According to the manufactureŕs recommendations, SsoFast EvaGreen Supermix was used to perform qPCR in a CFX96 Real-Time PCR System (BioRad). The amplification protocol started with a 30 s initial denaturation at 95°C and was followed by 45 cycles of 5 s at 95°C and 55°C for 10 s. Single product amplification was checked by recording the melting curves. For each transcript, dilution series of pooled cDNA were performed under the same conditions for primer efficiency calculations. *IPCS2* was amplified using primers JM231 and JM280 (Supplemental Table 1). *IPCS1* was amplified using the published primers by Ito et al. (2021). *IPCS1* and *IPCS2* expression levels were normalized to *ACTIN8* (amplified with primers EP223 and EP224; Supplemental Table 1).

### Yeast complementation assay

The premature stop codon insertion (W205*) or deletion of the remaining amino acids following W205 in IPCS2 were inserted in the previously published p426-*AtIPCS2* expression plasmid (Mina et al. 2010; Haslam et al. 2024) using the Q5 Site-Directed Mutagenesis Kit (New England Biolabs). All primers used for cloning are listed in Supplemental Table S1. The plasmid sequences were confirmed by Sanger sequencing. The previously reported LiAc/SS carrier DNA/PEG method was used to transform the different expression plasmids in parallel into *YPH499-HIS3-GAL1_pro_:AUR1* cell aliquots (Gietz and Schiestl 2007). Uptake of the correct plasmid was verified by colony PCR and Sanger sequencing for all colonies used for the complementation assay. Yeast strains were cultivated in or on synthetic complete dropout medium (Gietz and Woods 2002) with(out) the appropriate nutrients and carbohydrate supplementation.

### Sphingolipidomics

Microsomes were enriched from 10 mg of lyophilized Arabidopsis rosette tissue by following a previously described protocol (Abas and Luschnig 2010). The established monophasic extraction method using isopropanol:hexane:water (60:26:14, v/v/v) (Markham et al., 2006) was performed for lipid extraction. Half of the total lipid extract volume was taken for methylamine treatment to hydrolyze glycerolipids. This allowed a cleaner background by removing false positive signals from the sphingolipid analysis and was based on the protocol described by Markham (2013). The sphingolipid analysis followed the established protocol by Herrfurth et al. (2021). In short, methylamine-treated sphingolipids were separated by an ACQUITY UPLC system (Waters Corp.) supplied with an HSS T3 silica-based reversed-phase C18 column (100 mm x 1 mm, 1.8 µL; Waters Corp.). Ionization was performed by a chip based nano-electrospray using TriVersa Nanomate (Advion BioScience) with 5 µm internal diameter nozzles. An AB Sciex 6500 QTRAP tandem mass spectrometer (AB Sciex) was used for analysis, which was operated in positive ionization mode of the nanomate and in multiple reaction monitoring (MRM) mode of the mass spectrometer. 2 µl sample was injected by an autosampler set at 18°C. The flow rate if the sample separation was 0.1 ml/min. Solvent A contained methanol-20 mM ammonium acetate (3:7, v/v) with 0.1 % acetic acid (v/v), while solvent B was composed of tetrahydrofuran:methanol:20 mM ammonium acetate (6:3:1, v/v/v) with 0.1 % acetic acid (v/v). Sphingolipids were separated by a linear solvent gradient, starting from 65% B for 2 min, which increased to 100% B in 8 min, holding for 2 min and re-equilibrating to the starting conditions in 4 min. For the targeted analysis, ceramides and glycosylceramides were detected as [M+H]^+^ (precursor ions) and by the dehydrated LCB fragments (product ions) whereas glycosyl inositol phosphorylceramides were detected as [M+NH_4_]^+^ (precursor ions) and by the ceramide backbone (product ions).

A 20% aliquot was used for methyl esterification by sulfuric acid-catalyzed methanolysis described in Haslam et al. (2024) for fatty acid methyl ester (FAME) normalization. Gas chromatography with flame ionization detection (GC-FID) was employed for total fatty acid quantification. For relative comparisons among genotypes, the LC-MS MRM peak areas of individual lipid species were normalized to the total fatty acid content of each sample.

### Confocal microscopy

Confocal microscopy was performed on a Leica TCS SP8 Falcon system equipped with HyD SMD detector. mVenus and mCitrine fluorescent protein was excited with a pulsed white light laser at 514 nm and fluorescence emission was recorded between 530 and 600 nm. The excitation wavelength for GFP was 488 nm and its fluorescence emission was detected between 500 and 540 nm. To exclude chloroplast autofluorescence signals from images, a fluorescence lifetime gate was set between 0.4 and 6 ns. The TGN marker SYP61-CFP was excited using a 405 nm Diode laser and fluorescence emission was captured between 460 and 500 nm. The Leica LASX 3.5.5 software or the LIF Projector plugin in Fiji (-ImageJ v1.49m) was used to export z-stacks (12 consecutive focal plane images with a distance of 1 µm) as maximum projections. Brightness/contrast was adjusted with Adobe Photoshop CS5.

### Time-course experiment of pathogen-induced PEN3 and callose accumulation

To evaluate the subcellular behavior of PEN3 in the *ipcs2_W205*_* mutant, this mutant was crossed with the published stable transgenic *pPEN3:PEN3-GFP* line (Stein et al. 2006). Homozygous F3 plants and the mother plant were inoculated with *Bgh* and analyzed between 12 hpi and 24 hpi via CLSM. 20 *Bgh* interaction sites per leaf from three different plants (n = 3) were evaluated for PEN3-GFP FA patterns. Those images showing a clear focal accumulation pattern were used for quantification. Fiji was used to draw the FA as a region of interest, which was then analyzed for the mean intensity and the area. Evaluation of callose accumulation was performed as previously described by Underwood et al. (2022). Briefly, three Col-0 and *ipcs2_W205*_* plants were inoculated with *Bgh*. 1 leaf per plant was cleared and stained with aniline blue (Lipka et al. 2005). 50 *Bgh* appressoria per leaf were analyzed for presence or absence of aniline blue-stained callose.

### Protein work

Total protein extraction from Arabidopsis leaves and Western blot analysis were performed according to a previously published protocol (Petutschnig et al. 2010). The signal intensity and sensitivity of the anti-GFP (ChromoTek) antibodies was increased using the SuperSignal^TM^ Western Blot Enhancer Kit (Thermo Fisher Scientific) following the manufactureŕs instructions.

### Statistical Analysis

Statistical difference between different genotypes was calculated via one or two-way ANOVA (p < 0.05) or a two-tailed t-test (p < 0.05).

### Accession number

Sequence data for the genes mentioned in this article can be found in the TAIR data libraries under the following accession numbers: IPCS2, AT2G37940; IPCS1, AT3G54020; IPCS3, AT2G29525; EDS1, AT3G48090; PAD4, AT3G52430; SID2, AT1G74710.

## ACKNOWLEDGEMENTS

We thank Prof. Dr Ralph T. Schwarz and Dr. med vet. Hosam Shams-Eldin for kindly providing us the *YPH499-HIS3-GAL1_pro_:AUR1* yeast strain. Sincere thanks to Prof. William Underwood and Prof. Shunyuan Xiao for the *pPEN3:PEN3-GFP* line or the *eps* mutant, respectively. We are grateful for excellent technical support by Sabine Wolfarth, Sabine Freitag, Susanne Mester and Felicitas Glasenapp.

## AUTHOR CONTRIBUTIONS

V.L., I.F. and Y.B. supervised the project; J.M., V.L, T.M.H., C.H., I.F., N.E., Y.B. designed the research; J.M., T.M.H., C.H. and N.E. performed the experiments; J.M., T.M.H., C.H., I.F.and V.L. analyzed the data; J.M. wrote the original manuscript with input from T.M.H. and V.L. V.L. edited the paper with input from all authors.

## DATA AVAILABILITY STATEMENT

The data presented in this study are available in the article or supplementary material. Correspondence and requests for materials should be addressed to.

## SUPPORTING INFORMATION

**Supplemental Figure 1:**
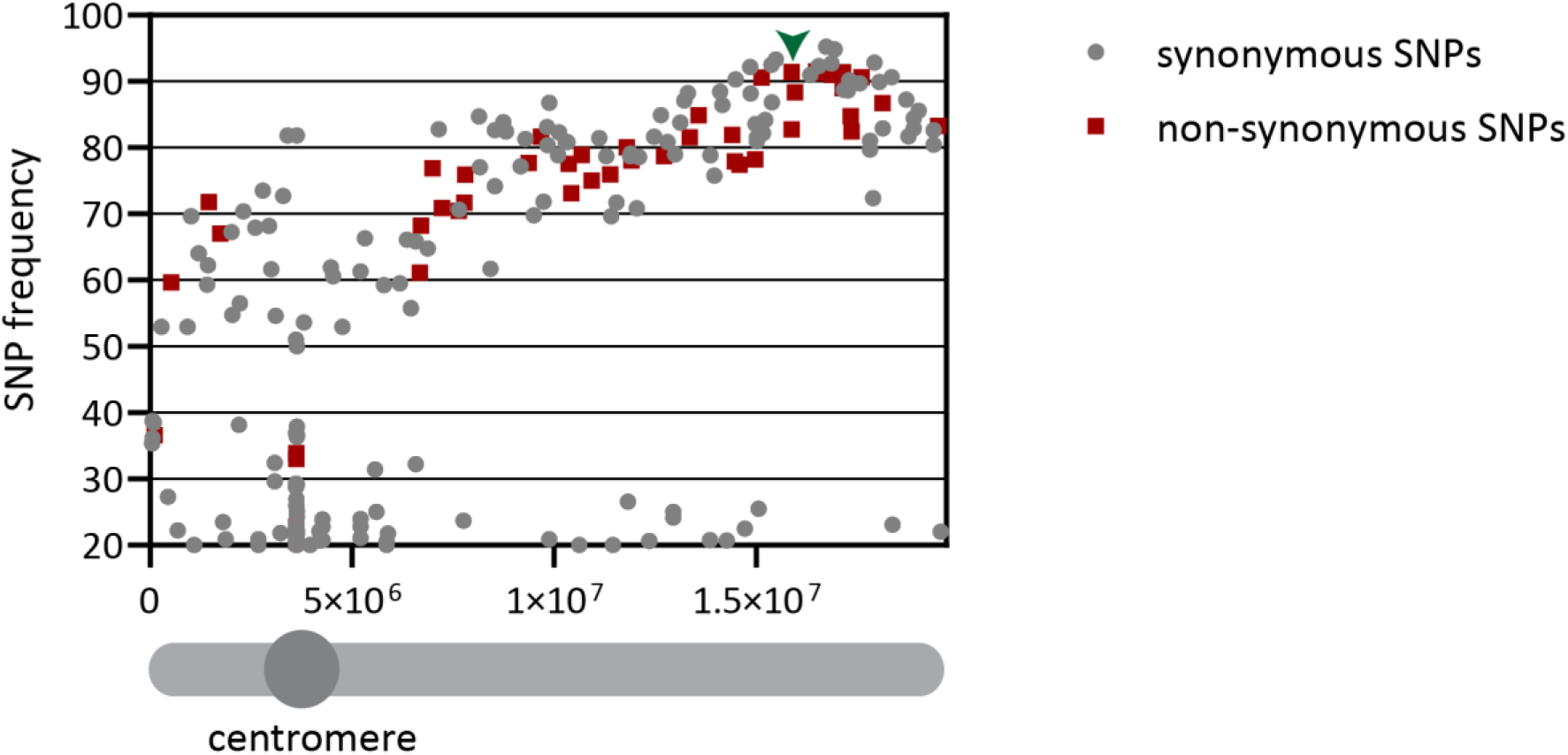
Mapping of *ipcs2_W205._* *ipcs2_W205*_* was backcrossed to Col-0 and the F2 generation was screened again for a *pen* phenotype. DNA of 28 plants with a clear *pen* phenotype was pooled and used for Illumina sequencing. After alignment of the reads to the TAIR10 genome, single nucleotide polymorphisms (SNPs) were called. The frequencies [%] of *ipcs2_W205*_* SNPs were plotted against their position [bp] on the respective chromosomes. This figure shows the frequency peak identified for chromosome 2. Synonymous mutations are shown as grey circles and non-synonymous mutations as red squares. The causative *ipcs2_W205*_* mutation in *AT2G37940* is indicated by a green arrowhead.

**Supplemental Figure 2:**
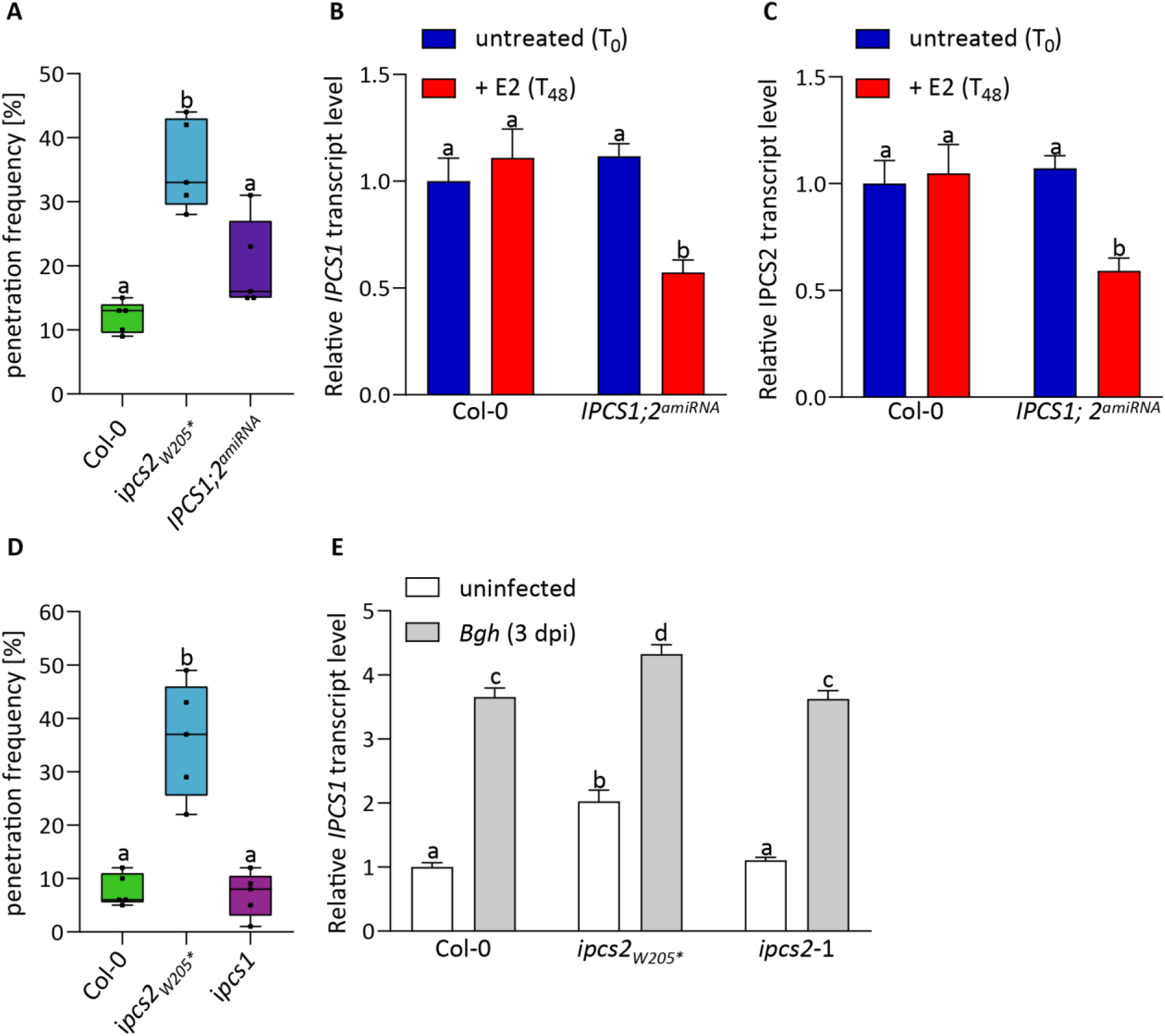
IPCS2 and not IPCS1 is the major contributor to Arabidopsis cell entry control against non-adapted powdery mildews. A: Col-0, *ipcs2_W205*_* and plants of the *IPCS1;2^amiRNA^*line were sprayed with β-Estradiol (E2) and 48 hours later infected with *Bgh*. Individual boxplots illustrate the penetration frequency of *Bgh* at 3 days post infection (dpi) and whiskers demonstrate minimum and maximum values. Letters show significant differences between genotypes (n =5; two-way ANOVA with Tukeýs post-hoc test; p < 0.05). The experiment was repeated twice with similar outcomes. B-C: Leaves of wild-type Col-0 and the *IPCS1;2^ariRNA^* line were harvested before (T_0_) and after E2-spraying (T_48_) and were analyzed by quantitative real-time PCR. Presented (B) *IPCS1* or (C) *IPCS2* transcript levels are relative to the untreated Col-0 sample. The bars represent the average ± standard error of the mean (n = 4-5), while letters indicate significant differences between genotypes and treatments (two-way ANOVA with Tukeýs post-hoc test; p < 0.05). Similar results were obtained in an additional independent experiment. D: *Bgh* penetration frequency on wild-type Col-0, *ipcs2_W205*_* and *ipcs1* mutant leaves at 3 dpi. Individual box plots include whiskers representing minimum and maximum values. Letters show significant differences between genotypes (n = 5; one-way ANOVA with Tukeýs post-hoc test; p<0.05). Similar results were confirmed in an additional independent experiment. E: Uninfected (white bars) or *Bgh*-infected (3 dpi) (grey bars) leaves of wild-type Col-0, *ipcs2_W205*_* and *ipcs2*-1 were used for quantitative real-time PCR analysis. Presented *IPCS1* transcript levels are relative to the uninfected Col-0 sample. The bars represent the average ± standard error of the mean (n = 4-5). Letters show significant differences between genotypes and treatments (two-way ANOVA with Tukeýs post-hoc test; p < 0.05). The experiment was repeated twice with similar outcomes.

**Supplemental Figure 3:**
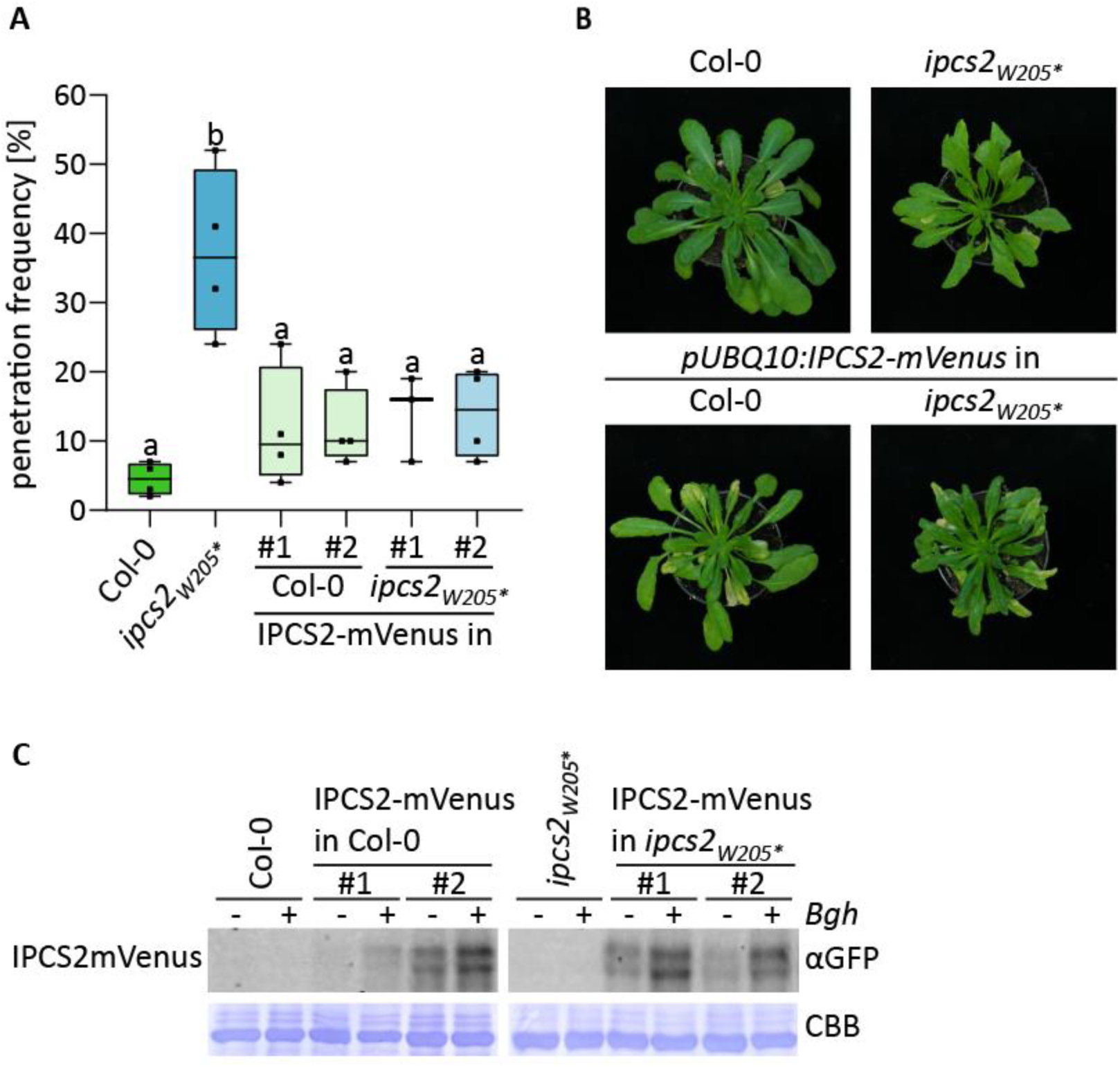
Complementation analyses of *ipcs2_W205_*. A: Col-0, *ipcs2_W205*_* and two independent transgenic plant lines stably expressing *pUBQ10:IPCS2-mVenus* in Col-0 or in the *ipcs2_W205*_* mutant background were infected with *Bgh*. Individual box plots represent the penetration frequency of *Bgh* at 3 days post infection. Whiskers show minimum and maximum values. Letters show significant differences between genotypes (n = 3-4; one-way ANOVA with Tukeýs post-hoc test; p<0.05). B: Macroscopic phenotype of Col-0, *ipcs2_W205*_* and the above described transgenic lines after 8-weeks under short-day conditions. C: Col-0, *ipcs2_W205*_* and the two above described independent transgenic lines expressing *pUBQ10:IPCS2-mVenus* in either Col-0 or *ipcs2_W205*_* were infected with *Bgh*. Leaf tissue of uninfected or *Bgh*-infected (24 hours post infection (hpi)) plants were used for total protein extraction and further Western Blot analysis using anti-GFP antibodies. Coomassie brilliant blue (CBB) staining was performed as a loading control. All results shown were replicated in an additional independent experiment.

**Supplemental Figure 4:**
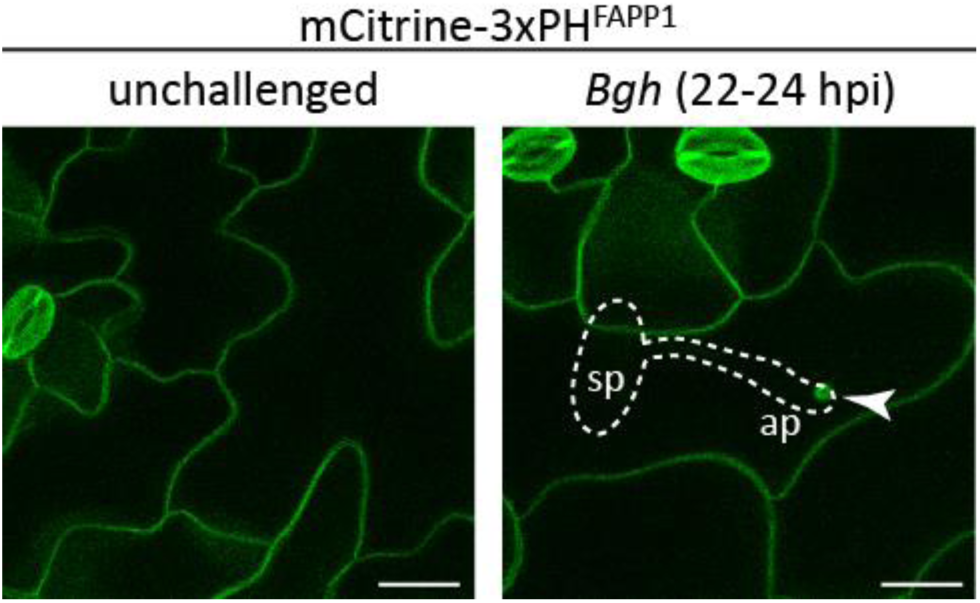
PI4P accumulates at *Bgh* contact sites. Maximum z-projections of CLSM images from unchallenged (left panel) or *Bgh*-infected (22-24 hours post infiltration (hpi); right panel) leaf epidermal cells of the PI4P biosensor marker line mCitrine-3xPH^FAPP1^. Arrowheads indicate *Bgh* penetration sites, while white lines illustrate fungal spore with appressorium formation. Similar results were obtained in an additional independent experiment. Ap = appressorium; sp = spore. Scale bar = 20 µm.

**Supplemental Figure 5:**
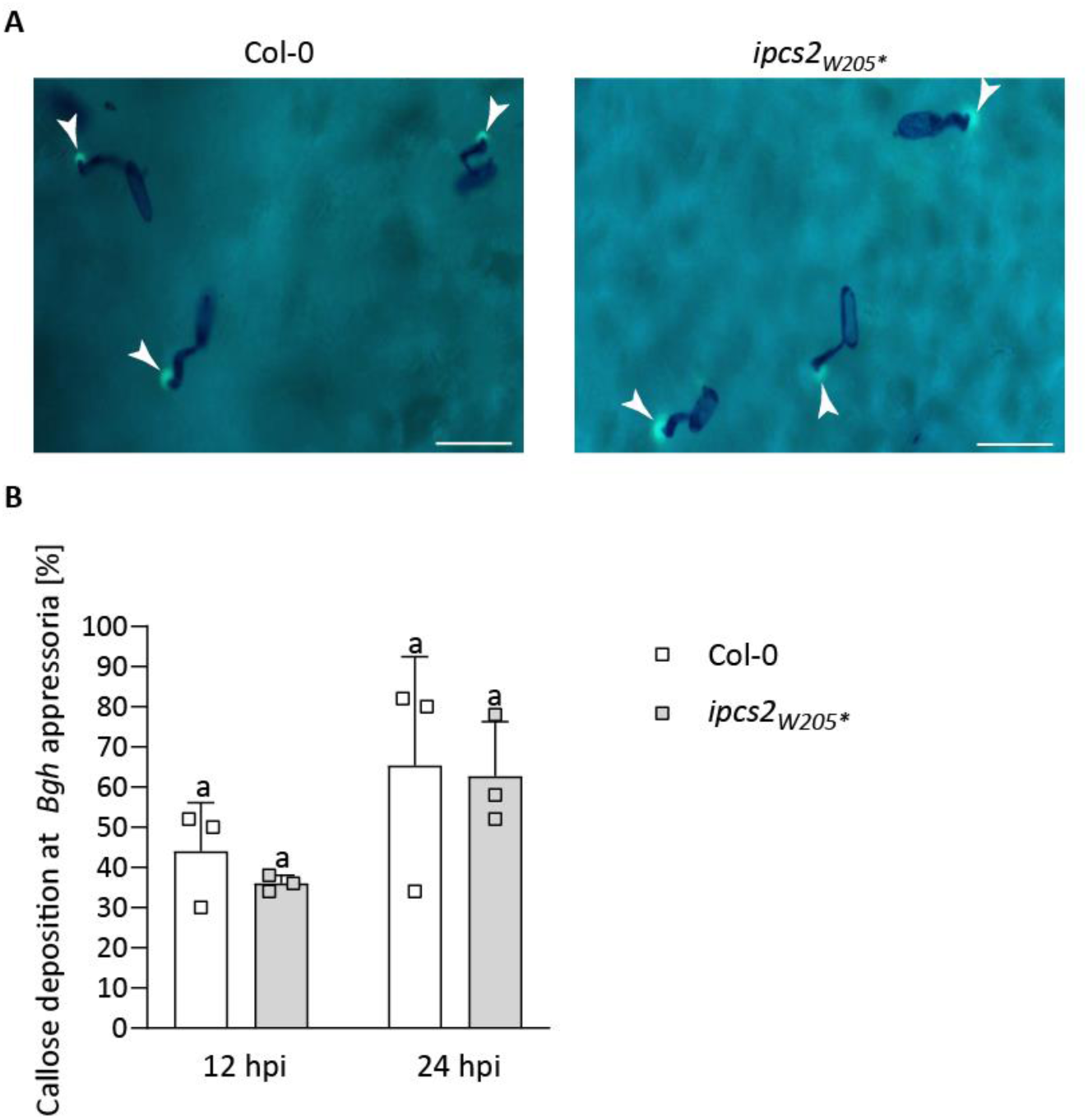
Callose deposition at *Bgh* penetration sites is IPCS2-independent. A: Micrographs of aniline-blue- and Coomassie-brilliant-blue-stained leaves of Col-0 (left panel) or *ipcs2_W205*_* (right panel) after infection with *Bgh* (24 hours post infection (hpi)). Arrowheads indicate callose-enriched papillae at *Bgh* penetration sites. Scale bar = 50 µm. B: Frequency of callose deposition at 50 *Bgh* appressoria at 12 hpi and 24 hpi in Col-0 (white bars) or *ipcs2_W205*_* background (grey bars). Bars represent averages of three biological replicates ± standard deviation. Letters indicate significance at p < 0.05 determined by one-way ANOVA with Tukey’s post-hoc test. Experiment was repeated thrice with similar outcomes.

**Supplemental Figure 6:**
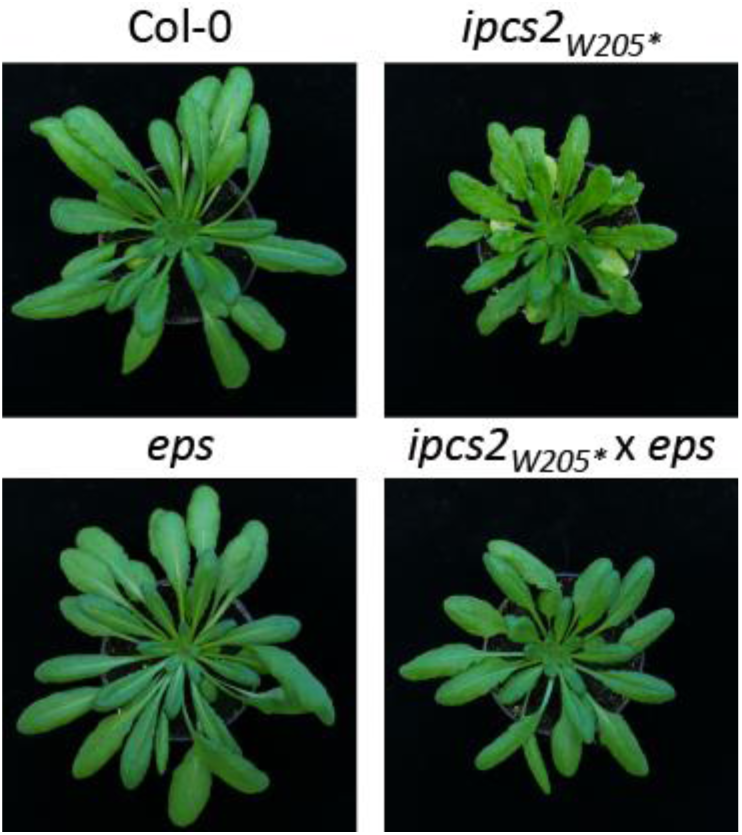
The spontaneous lesion phenotype of *ipcs2_W205*_* is SA-dependent. Macroscopic phenotype of Col-0, *ipcs2_W205*_*, *eps* and the *ipcs2_W205*_* x *eps* double mutant after 8-weeks under short-day conditions.

**Supplemental Table 1:**
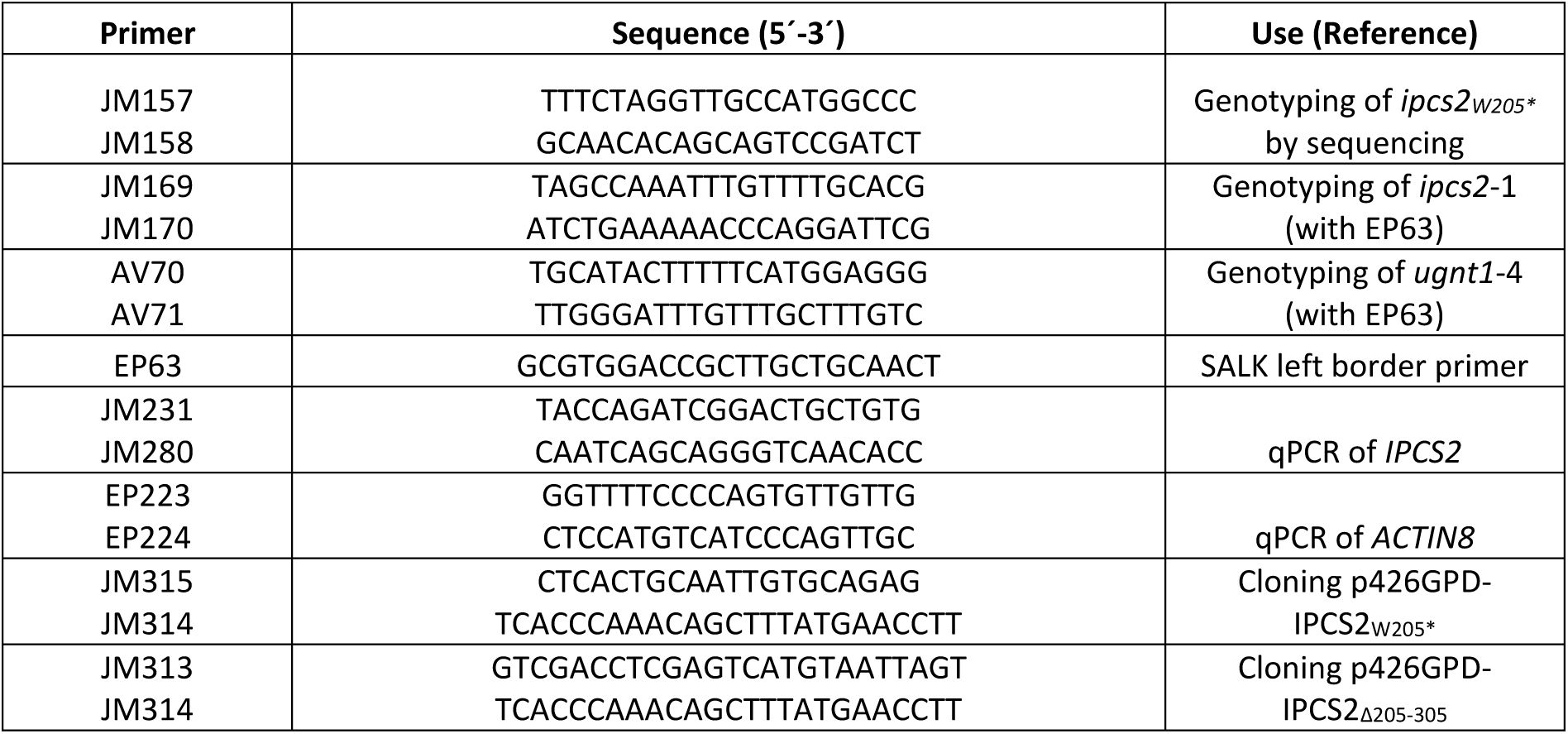
Oligonucleotides used in this study.

